# Chromosome level genome assembly and annotation of highly invasive Japanese stiltgrass (*Microstegium vimineum*)

**DOI:** 10.1101/2021.09.16.460655

**Authors:** Dhanushya Ramachandran, Cynthia D. Huebner, Mark Daly, Jasmine Haimovitz, Thomas Swale, Craig F. Barrett

## Abstract

The invasive Japanese stiltgrass (*Microstegium vimineum*) affects a wide range of ecosystems and threatens biodiversity across the eastern USA. However, the mechanisms underlying rapid adaptation, plasticity, and epigenetics in the invasive range are largely unknown. We present a chromosome-level assembly for *M*. *vimineum* to investigate genome dynamics, evolution, adaptation, and the genomics of phenotypic plasticity. We generated a 1.12 Gb genome with scaffold N50 length of 53.44 Mb respectively, taking a *de novo* assembly approach that combined PacBio and Dovetail Genomics Omni-C sequencing. The assembly contains 23 pseudochromosomes, representing 99.96% of the genome. BUSCO assessment indicated that 80.3% of Poales gene groups are present in the assembly. The genome is predicted to contain 39,604 protein-coding genes, of which 26,288 are functionally annotated. Furthermore, 66.68% of the genome is repetitive, of which unclassified (35.63%) and long terminal repeat (LTR) retrotransposons (26.90%) are predominant. Similar to other grasses, *Gypsy* (41.07%) and *Copia* (32%) are the most abundant LTR-retrotransposon families. The majority of LTR-retrotransposons are derived from a significant expansion in the past 1-2 million years, suggesting the presence of relatively young LTR-retrotransposon lineages. We find corroborating evidence from Ks plots for a stiltgrass-specific duplication event, distinct from the more ancient grass-specific duplication event. The assembly and annotation of *M*. *vimineum* will serve as an essential genomic resource facilitating studies of the invasion process, the history and consequences of polyploidy in grasses, and provides a crucial tool for natural resource managers.

**Significance:** The current lack of genomic resources for the invasive Japanese stiltgrass—and thousands of other invasive species globally—severely limits our understanding of the invasion process and hinders decision-making for effective management and control. In this study, we present a chromosome-level genome assembly and annotation of Japanese stiltgrass, a problematic weed in eastern North America, identifying a clear history of polyploidy and recent activity of transposable elements. The ultimate goal is to advance genomic studies to better understand the dynamics of non-native species during the various invasion phases, thereby providing insights into effective control strategies to manage current and future invasions.

## Introduction

Invasive species cause billions of dollars in damage annually, and are considered the second greatest threat to native biodiversity after habitat loss (Lechar and Mooney, 2009; Simberloff, 2013). Yet, genomic resources for invasive species are generally lacking relative to other economically important species such as crops, microbial pathogens, and many animal systems (McCartney, Mallez, & Gohl, 2019). Almost half of the native species in the United States are at risk of extinction either due the direct effects of introduced species or impacts combined with other processes (Pimentel et al. 2005). Efforts to identify and eradicate newly introduced species are hampered by the lack of resources needed to predict how and why some species will become invasive. Genomics has become an increasingly valuable and cost-efficient tool to predict and diagnose invasions (Chown et al. 2015; Hamelin and Roe, 2020). Genomics can provide novel insights on the roles of genetic variation, multiple introductions, admixture, introgression, and rapid adaptation (Schrader et al. 2014, Malinsky 2019, Kreiner et al, 2019, Bertolotti et al. 2020, Olazcuaga et al., 2020, Yainna et al. 2020). For instance, a high-quality genome is useful for genome-wide scans of selection, trait association mapping, and timing invasion events (e.g. North et al. 2021, DeGiorgio et al., 2016; Nielsen et al., 2005). With improved understanding and forecasting at each stage of the invasion process, managers can make decisions on invasions much more accurately than in the past (Bergeron et al. 2019, Keriö et al. 2020). Hence, sequencing whole genomes for these non-model organisms provide crucial tools to efficiently manage and predict future invasions.

Japanese stiltgrass (*Microstegium vimineum*) is a shade-tolerant, annual, C4 grass introduced to the eastern USA from Asia in the early 1900s that has spread to 30 US states and Canada. This species invades a range of habitats in the USA, displays a high degree of phenotypic plasticity, has a mixed mating system (outcrossing and self-fertilization), and exhibits prolific reproductive output with seeds being viable in the soil up to five years (Barden 1987, Gibson et al. 2002, Redman 1995, Nees 2016, Culpepper et al. 2018). Considerable research interest has been focused on unraveling potential links between ploidy levels and invasiveness, as most invasive plant species are polyploids (Pandit et al 2011, te Beest et al 2012). Japanese stiltgrass is an ideal system for the study of rapid adaptation of invasive species, being a putative polyploid in addition to the aforementioned features (2N = 20 as opposed to the ‘base’ 2N = 10 among members of Andropogoneae; Watson and Dallwitz, 1992).

Here, we present a high-quality, chromosome-level assembly and annotation for *M*. *vimineum*, by integrating PacBio sequencing, Omni-C scaffolding, and RNAseq. The genome will lay groundwork for further investigation of traits allowing *M*. *vimineum* to adapt and thrive as an invasive species. Further, this genome will provide an important genomic resource for studies of rapid adaptation in invasive plants, help elucidate the history and consequences of polyploidy in grasses, and provide a tool for natural resource scientists and managers.

## Materials and Methods

### Sample collection and DNA extraction

Florets containing seeds were collected and mixed from three populations in the Potomac Ranger District (PRD) of the Monongahela National Forest (MNF) near Petersburg, West Virginia, USA, and three populations in the Cheat Ranger District (CRD) of the same forest near Parsons, West Virginia, USA. Florets were air-dried for 3 months, and cold-dry stratified at 4°C for one year. One plant was also grown from seed-bank soil collected along the Monongahela River Rail Trail (RT) in Morgantown, West Virginia, USA. Seeds were germinated over two weeks in a Conviron growth chamber under temperatures of 25°C/15°C (12-hour day/12-hour night), approximately 70% humidity, and 500 μmol m^−2^ s^−1^ light. RT seedlings were transplanted into potting soil. After germination, day length was increased to 14 hours and night temperature was increased to 20°C. The complete shoot of one individual was harvested from each location (PRD, CRD, and RT). Twenty-five grams of fresh, young, green leaf tissue from one PRD accession was chosen for genome sequencing; tissue was flash-frozen in liquid nitrogen and stored at −80°C for one month before shipping on dry ice. The remainder of these individuals were stored at −80°C upon flowering with a voucher specimen of each deposited at the Northern Research Station, USDA Forest Service Herbarium. Further, tissue was harvested from these frozen samples for RNA-seq analysis. Approximately 0.2g of tissue was harvested from you, developing tissues for: 1) leaves, 2) roots, 3) cleistogamous inflorescences (covered by leaf sheaths at the nodes), and 4) apical, chasmogamous inflorescences. Tissues were flash frozen as above, stored at −80°C, and shipped on dry ice to GeneWiz, Inc. (South Plainfield, New Jersey, USA) for RNA sequencing.

### PacBio Library Sequencing

Total genomic DNAs were extracted from leaf tissues to construct sequencing libraries (Supplementary Material). PacBio SMRTbell libraries (∼20kb) were constructed using the SMRTbell Express Template Prep Kit 2.0 (PacBio, Menlo Park, CA, USA), following the manufacturer’s protocol. Libraries were bound to polymerase using the Sequel II Binding Kit 2.0 (PacBio) and loaded onto a PacBio Sequel II at Dovetail Genomics, LLC. Sequencing was performed on two PacBio Sequel II 8M SMRT cells. PacBio reads were assembled using the Wtdbg2 pipeline (Ruan and Li, 2020). Contaminants and ‘haplotigs’ (contigs from a single, alternative haplotype) were filtered using Blobtools v1.1.1 (Laetsch and Blaxter, 2017) and purge_dups v1.1.2 (Guan et al. 2019; Supplementary Material).

### Dovetail Omni-C Library Preparation and Sequencing

For Dovetail Omni-C libraries, chromatin was fixed with formaldehyde, extracted, and randomly digested with DNAse I. Chromatin ends were repaired and ligated to a biotinylated bridge adapter, followed by proximity-ligation of adapter-containing ends. After proximity ligation, crosslinks were reversed and DNA was purified. Purified DNA was treated to remove biotin that was not internal to ligated fragments, and sequencing libraries were generated using NEBNext Ultra enzymes and Illumina-compatible adapters. Biotin-containing fragments were isolated using streptavidin beads before PCR enrichment of each library. The library was sequenced on an Illumina HiSeqX platform to produce approximately 30× sequence coverage depth. HiRise was used for scaffolding, a pipeline designed specifically for proximity ligation data (Putnam et al, 2016), requiring mapping quality (MQ) > 50 reads. Dovetail OmniC library sequences were aligned to the draft input assembly using bwa (https://github.com/lh3/bwa). Separations of Dovetail OmniC read pairs mapped within draft scaffolds were analyzed by HiRise to produce a likelihood model for genomic distance between read pairs, and used to identify and break putative mis-joins, to score and make prospective joins.

### RNA-seq

Total RNAs were extracted using the QIAGEN RNeasy Plus Kit following manufacturer protocols. Total RNAs were quantified using the Qubit RNA Assay and a TapeStation 4200. Prior to library preparation, DNase treatment was performed followed by AMPure (Beckman Coulter Life sciences) bead cleanup and QIAGEN FastSelect HMR rRNA (QIAGEN) depletion. Libraries were prepared with the NEBNext Ultra II RNA Library Prep Kit following manufacturer protocols and run on an Illumina NovaSeq6000 in 2×150 bp configuration.

### Assessment of genome assembly quality

Completeness of the genome and predicted gene quality was assessed using Benchmarking Universal Single-Copy Orthologs (BUSCO v3.0.1; Simao et al. 2015). The poales_odb10 lineage-specific profile that contains 4,896 BUSCO gene groups was evaluated against our chromosome-level assembly.

### Gene prediction and annotation

Coding sequences from *Coix lacryma*-*jobi* (PRJNA544872), *Miscanthus sacchariflorus* (PRJNA435476), *Saccharum ‘*hybrid cultivar’ (PRJNA272769), *Sorghum bicolor* (PRJNA331825), and *Zea mays* (PRJNA10769) were used to train the *ab initio* model for *Microstegium vimineum* using AUGUSTUS (version 2.5.5; Stanke et al. 2008). The same coding sequences were also used to train a separate *ab initio* model for *Microstegium vimineum* using SNAP (v2006-07-28; Korf, 2004). RNA-seq reads were mapped to the genome using STAR (v2.7; Dobin et al. 2013) and intron-exon boundary hints were generated. AUGUSTUS was then used to predict genes in the repeat-masked reference genome. Only genes predicted by both SNAP and AUGUSTUS were retained in the final gene sets. To assess quality of gene predictions, AED (Annotation Edit Distance) scores were generated for predicted genes in MAKER. Genes were further characterized for putative functions by performing a BLAST search of peptide sequences against the UniProt database. tRNAs were predicted using the software tRNAscan-SE (version 2.05, Chan & Lowe 2019). More details on gene prediction and annotation is described in the Supplementary Material.

### Repeat analysis

Repeat families in *Microstegium vimineum* were identified *de novo* and classified using RepeatModeler (version 2.0.1; Flynn et al. 2020) and EDTA v1.9.4 (Ou et al. 2019). RepeatModeler uses RECON (version 1.08; Bao and Eddy, 2002) and RepeatScout (version 1.0.6; Price et al. 2005) for *de novo* identification. Class I LTR-retrotransposons (LTR-RT) were further predicted and annotated using RepeatModeler and EDTA. Both tools use a series of LTR-RT identification programs such as LTR-harvest, LTR-finder, and LTR-retriever. Redundant and nested insertions were removed by EDTA. Intact LTR-RTs were identified and approximate insertion times (million years ago) were estimated using LTR-retriever (based on a grass-specific LTR substitution rate of 1.3 × 10^−8^ mutations/site/year; Ma and Bennetzen, 2004). EDTA further uses TIR-learner and Helitron-scanner to predict and annotate Class II DNA transposons and helitrons, or rolling circle DNA transposons (Kapitonov and Jurka, 2001, Feschotte and Wessler, 2001). The custom repeat library obtained from RepeatModeler and EDTA was used to discover, identify, and mask repeats in the assembly using RepeatMasker (Version 4.1.0; http://www.repeatmasker.org).

### Detection of whole genome duplication events

To investigate whole genome duplication (WGD) events in *M*. *vimineum* genome, the distribution of synonymous substitution (Ks) rates was obtained from protein coding sequences and compared with closely related grasses, e.g. *Sorghum bicolor*, *Coix lacryma*-*jobi*, and *Zea mays*. Paralog and ortholog pairs were detected from protein sequence data and the associated Ks values were calculated using the tool “ksrates” (https://github.com/VIB-PSB/ksrates; Sensalari et al. 2021). A mixed Ks plot was generated by comparing ortholog-Ks estimates to the paralog-Ks scale of *M*. *vimineum*. MCScan (https://github.com/tanghaibao/jcvi/wiki/MCscan-(Python-version); Tang et al. 2008) was used for pairwise synteny (protein) search with the LSAT results of *M*. *vimineum* versus *S*. *bicolor*. The MCScan “jcvi.graphics.dotplot” module was used to visualize pairwise synteny results. Further, the genes of *M*. *vimineum* genome were classified into singletons, dispersed, tandem, proximal, and WGD/segmental duplicates using ‘duplicate_gene_classifier’ module within the MCScan_X tool (Wang et al. 2012), by parsing the all_Vs_all BLASTp results.

## Results and Discussion

### Genome sequencing and assembly

We generated a high-quality, chromosome-level genome assembly of *M*. *vimineum* using PacBio and Dovetail Omni-C libraries. Using approximately 60 Gb of PacBio long read data, we initially assembled 5,261 *de novo* contigs with N50 of 605 kb. In parallel, a total of 73.21 Gb (30× coverage) of short read sequence data were produced by Illumina HiSeqX from Dovetail’s Omni-C libraries to achieve chromosome-scale scaffolding. The initial assembly was significantly improved with Omni-C data using the HiRise pipeline (Fig. 1A), which produced a final assembly consisting of 462 scaffolds spanning 1.1 Gb in length, with the scaffold N50 size of 53 Mb (Table 1). The final assembly covers 99.96% of 1.3 Gb genome size and interestingly, about 99.11% of assembled genome were anchored into 23 pseudochromosomes (size range 20.9-68.32 Mb), corresponding closely to the expected number of 20 chromosomes (Fig 1A).

**Table 1.**
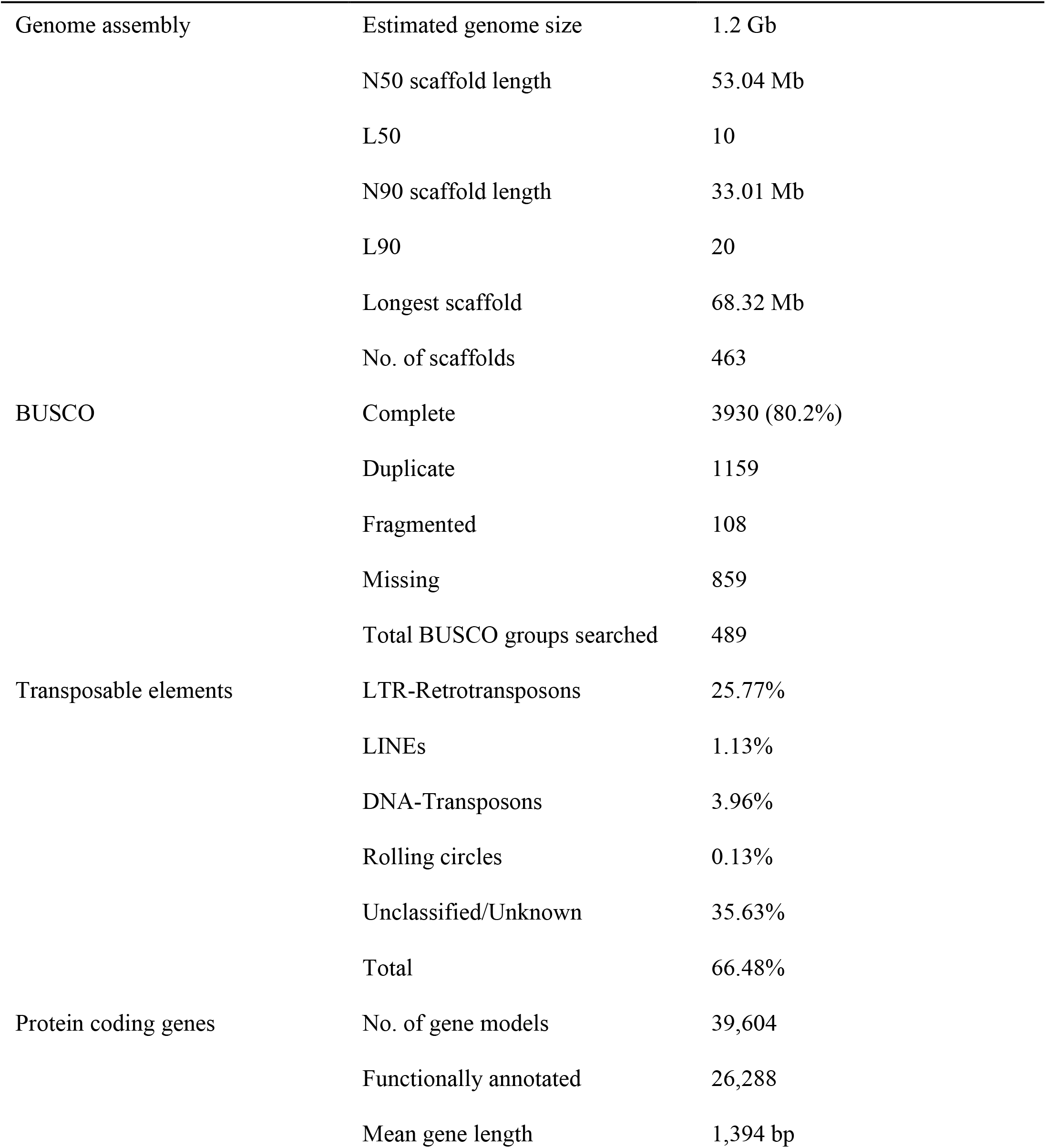
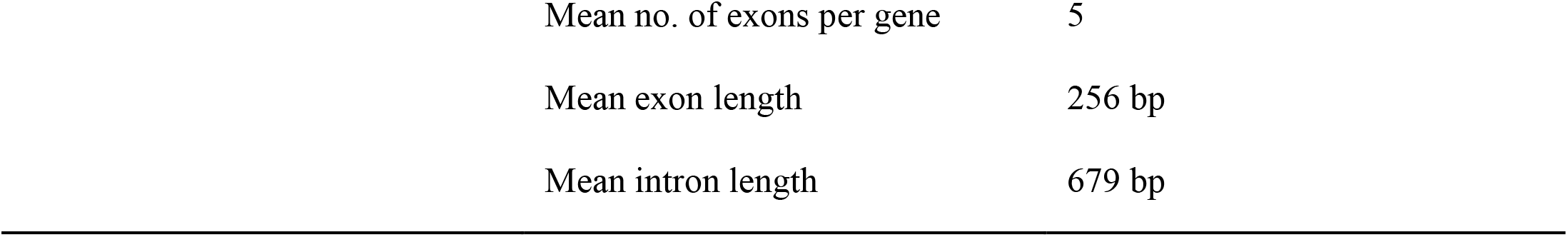
Summary of the genome assembly and annotation

|  |  |  |
| --- | --- | --- |
| Genome assembly | Estimated genome size | 1.2 Gb |
|  | N50 scaffold length | 53.04 Mb |
|  | L50 | 10 |
|  | N90 scaffold length | 33.01 Mb |
|  | L90 | 20 |
|  | Longest scaffold | 68.32 Mb |
|  | No. of scaffolds | 463 |
| BUSCO | Complete | 3930 (80.2%) |
|  | Duplicate | 1159 |
|  | Fragmented | 108 |
|  | Missing | 859 |
|  | Total BUSCO groups searched | 489 |
| Transposable elements | LTR-Retrotransposons | 25.77% |
|  | LINEs | 1.13% |
|  | DNA-Transposons | 3.96% |
|  | Rolling circles | 0.13% |
|  | Unclassified/Unknown | 35.63% |
|  | Total | 66.48% |
| Protein coding genes | No. of gene models | 39,604 |
|  | Functionally annotated | 26,288 |
|  | Mean gene length | 1,394 bp |

|  |  |
| --- | --- |
| Mean no. of exons per gene | 5 |
| Mean exon length | 256 bp |
| Mean intron length | 679 bp |

**Fig. 1.**
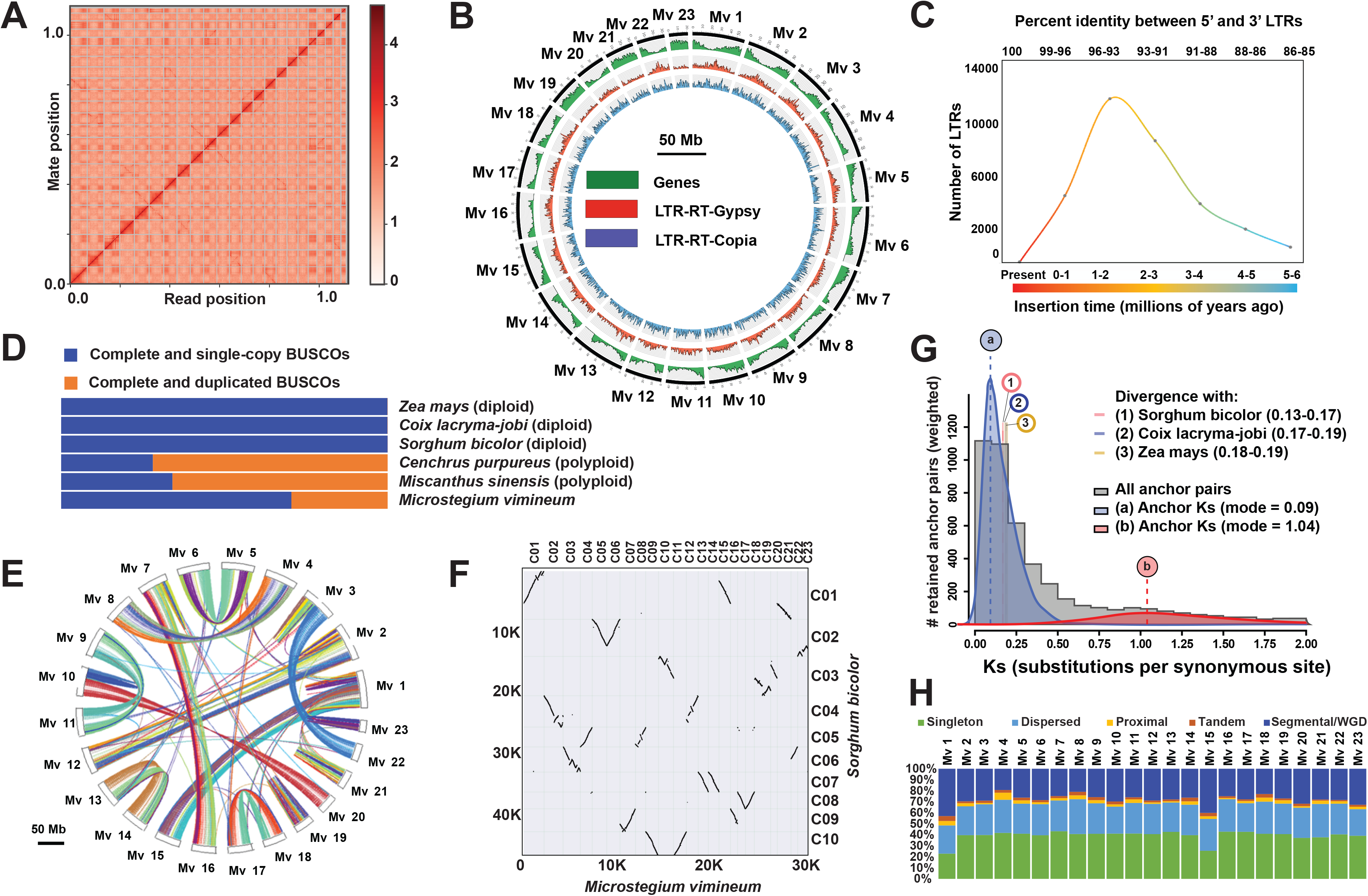
**A**. Linkage density heatmap of the *Microstegium vimineum* genome. The x and y axes represent the mapping positions of the first and second read in a read pair, respectively. The diagonal lines from lower left to upper right in the plot represent each of the 23 *M*. *vimineum* pseudochromosomes. Dots (sequences) outside the diagonal are likely repetitive sequences that occur in multiple chromosomes. **B**. Circos plot of *M*. *vimineum* genome assembly showing distributions of genes (green), *Gypsy* LTR-RTs (red), and *Copia* LTR-RTs (blue). **C**. Insertion age estimates of LTR-retrotransposons in million years ago (mya) based on a grass-specific LTR mutation rate (Ma and Bennetzen, 2004). **D**. BUSCO assessment results of orthologs among *M*. *vimineum*, closely related diploids (*Sorghum bicolor*, *Coix lacryma*-*jobi*, *Zea mays*), and polyploids (*Miscanthus sinensis* and *Cenchrus purpureus)*. **E**. Inter-chromosomal synteny with links representing syntenic blocks between *M*. *vimineum* chromosomes. **F**. Macrosynteny dotplot of *M*. *vimineum* and *Sorghum bicolor* chromosomes displaying large-scale duplications, inversions, and translocations. **G.** The frequency distributions of synonymous substitution rates (Ks) of homologous gene pairs located in the collinearity blocks of *M*. *vimineum*. The Ks distribution for *M*. *vimineum* is shown in gray, with two WGD peaks indicated in blue and red. The vertical lines labeled ‘a’ and ‘b’ indicate the modes of these peaks, which are taken as Ks-based WGD age estimates. The numbered vertical lines represent rate-adjusted mode estimates of one-to-one ortholog Ks distributions between *M*. *vimineum* and closely related species, representing speciation events. **H**. Distributions of gene duplicate origins across each chromosome in *M*. *vimineum* genome.

### Repeat and gene annotation

Over half of the genome is composed of repetitive elements (66.68%, 745.92 Mb; Table 1). Class I LTR retrotransposons are predominant, constituting 39.08% of the assembled genome. Similar to other grasses, the most abundant LTR-retrotransposon family present in *M*. *vimineum* genome is *Gypsy* (41.07%), followed by *Copia* (32%) (Baucom et al. 2009, Schnable et al. 2009, Tian et al. 2009, Paterson et al. 2009). *Gypsy* elements are distributed in gene-poor regions in most pseudochromosomes, while *Copia* shows a more even distribution (Fig 1B). Calibrated sequence divergence of 5’ and 3’ terminal repeats revealed that most LTR-retrotransposons insertions appear to have occurred 1-2 mya (Fig 1C), suggesting recent activity of LTR-retrotransposons and preponderance of young LTR lineages in the genome.

We predicted 39,604 genes spanning 55.22 Mb (approximately 4.9%) of the genome, with an average gene length of 1,394 bp (Table 1). A total of 26,230 genes were functionally annotated. We evaluated the completeness of the predicted gene sets and extent of gene duplication with 4,896 BUSCOs from the Poales database (v10; Manni et al. 2021), of which 3,930 (80.2%) were complete, indicating a relatively complete genome assembly and gene prediction (Table 1). An interesting observation among the complete BUSCO’s was the presence of 1,159 (30%) complete duplicated copies. This degree of duplication is comparable with, but lower than that seen in the polyploids *Miscanthus sinensis* (Mitros et al. 2020) and *Cenchrus purpureus* (Yan et al. 2021) (Fig 1D).

### Whole genome duplication in *M*. *vimineum*

Syntenic blocks in *M*. *vimineum* are displayed in Fig. 1E. Investigation of collinear orthologs between *M*. *vimineum* and the diploid *Sorghum bicolor* revealed a 2:1 (*M. vimineum:S. bicolor*) synteny pattern with evidence of duplications, translocations, and inversions confirming the occurrence of whole genome duplication in *M*. *vimineum*. Chromosomes 13 and 14 in *M*. *vimineum* are apparent homeologs, displaying collinearity along their entire length to *Sorghum* chromosome 7. Large-scale inversions are observed on *M*. *vimineum* chromosomes 5 and 6, which are syntenic to *Sorghum* chromosome 2. Inverted homeologs within chromosomes 17 and 18 of *M*. *vimineum* display clear collinearity to *Sorghum* chromosome 8. Chromosomes 4 and 8 of *M*. *vimineum* are syntenic to *Sorghum* chromosome 5, but with two large-scale inversions in *M*. *vimineum* chromosome 8 (Fig 1F).

The Ks peaks in Fig. 1G indicate two WGD events: 1) a paleoduplication event shared by all grasses at Ks=1.04, estimated at 80–90 million years ago (mya; Paterson et al. 2004), and 2), and a *M*. *vimineum*-specific WGD at Ks=0.09. The majority of duplicates in *M*. *vimineum* were derived from WGD/segmental (28.5%) and dispersed (27.5%) duplications, corroborating polyploidization followed by considerable chromosomal reshuffling in *M*. *vimineum* (Fig 1H). At a minimum, this suggests the *M*. *vimineum*-specific duplication likely occurred in the last ∼10mya, but additional taxon sampling is needed to more accurately estimate the timing of this event.

## Conclusion

We generated a high-quality, chromosome-scale genome assembly and annotation for *Microstegium vimineum* using PacBio sequencing and Omni-C technology. Genome quality assessment indicated a highly contiguous, accurate assembly and annotation, revealing recent WGD and transposon activity. Given the paucity of sequenced genomes for invasive species, this genome will serve as an important resource to study invasive species at the genomic level. Due to the varying abilities of introduced species to establish in a new environment, decision-making regarding resource allocation, mitigation, and management has always been uncertain; availability of genomic information for non-native species may provide new solutions (Hamelin and Roe, 2019). Whole-genome information expedites downstream population genomic studies on the role of multiple introductions, admixture, and adaptive ramifications of novel genotypes allowing ‘exploration’ of novel phenotypic space, phenologies, and ecological interactions (Bertolotti et al., 2020). Also, this genome will facilitate studies on the role of epigenetic variation and mobile elements of the genome to delineate their roles in rapid adaptation to the introduced range. These latter processes may allow novel phenotypes and gene expression modifications against the predicted genomic background of low allelic diversity in many invasive species (Mérel et al., 2021). Further, comparative genomics and evolutionary studies of invasive versus non-invasive grasses or other plants, animals, and microbes may help to identify genomic commonalities characteristic of successful invaders.

## Acknowledgments

Funding was provided by National Science Foundation Award OIA-1920858 and a Dovetail Genomics Tree of Life Grant to CFB. Growth chamber access was provided by the US Department of Agriculture Forest Service (Northern Research Station, Morgantown, West Virginia, USA). We thank Joanna Gallagher, and Jordan Zhang at Dovetail Genomics, LLC, for assistance with genome sequencing and assembly, and Michael McKain for helpful discussion and assistance with data analyses.

## Data Availability

Sequence Read Archive: PacBio, Omni-C, and RNAseq – BioProject and BioSamples GenBank: Genome and annotation

## Supplementary Material

### DNA extraction for PacBio Library preparation

Total genomic DNAs were extracted using the CTAB method (Doyle and Doyle, 1987). DNA concentration was quantified with a Qubit Fluorometer v2.0 using the dsDNA Broad Range assay (Thermo Fisher Scientific, Waltham, Massachusetts, USA), yielding a total of 11.2μg. Genomic DNA was further purified using a Qiagen Mini Column, following the manufacturer’s protocol (Qiagen, Hilden, Germany). The final DNA sample was resuspended in 50μl of Tris-EDTA Buffer (pH 8.0). The sample was quantified and checked for high molecular weight via Qubit, NanoDrop (Thermo Fisher Scientific, Waltham, Massachusetts, USA), Pulse Field Gel Electrophoresis, and an Agilent TapeStation 4200 (Santa Clara, California, USA).

### PacBio reads assembly using Wtdbg2 Assembler and Scaffolding with Dovetail Omni-C

Wtdbg2 (Ruan and Li, 2020) was run with the following parameters: −x sq −g 1g −L 5000. Blobtools v1.1.1 (Laetsch and Blaxter, 2017) was used to identify potential contamination in the assembly based on blast (v2.9) results of the assembly against the NT database. A fraction of the scaffolds was identified as contaminant and were removed from the assembly. The filtered assembly (filtered.asm.cns.fa) was then used as an input to purge_dups v1.1.2 (Guan et al. 2019) and potential haplotypic duplications were removed from the assembly, resulting in the final purged.fa assembly. Contig-contig read pair linkages are analyzed to produce a 3D model of the genome. This model accounts for the distance between contigs and number of supporting Omni-C read pairs between each contig. The HiRise software was then used ranks contigs based on these linkages and scaffolds were scored based on observed insertion size and evidence inside these insertion size windows (Putnam et al, 2016).

### Gene prediction and annotation

Coding sequences from *Coix lacryma*-*jobi* (PRJNA544872), *Miscanthus sacchariflorus* (PRJNA435476), *Saccharum ‘*hybrid cultivar’ (PRJNA272769), *Sorghum bicolor* (PRJNA331825), and *Zea mays* (PRJNA10769) were used to train the ab initio model for *Microstegium vimineum* using AUGUSTUS (version 2.5.5; Stanke et al. 2008). Six rounds of prediction optimization were done with AUGUSTUS. The same coding sequences were also used to train a separate ab initio model for *Microstegium vimineum* using SNAP (v2006-07-28; Korf, 2004). RNA-seq reads were mapped to the genome using STAR (v2.7; Dobin et al. 2013) and intron hits generated with the ‘bam2hints’ tools within AUGUSTUS. MAKER, SNAP and AUGUSTUS (with intron-exon boundary hints provided from RNA-Seq) were then used to predict for genes in the repeat-masked reference genome. To help guide the prediction process, Swiss-Prot peptide sequences from the UniProt database (The UniProt Consortium, Nucleic Acids Research, 2021) were downloaded and used in conjunction with the protein sequences from the species above to generate peptide evidence in the MAKER pipeline (Cantarel et al. 2008). Only genes that were predicted by both SNAP and AUGUSTUS were retained in the final gene sets. To help assess the quality of the gene prediction, AED scores were generated for each of the predicted genes in MAKER. Genes were further characterized for putative functions by performing a BLAST search of peptide sequences against the UniProt database. tRNAs were predicted using the software tRNAscan-SE (version 2.05, Chan & Lowe 2019).

## Literature cited

Bao Z, Eddy SR. 2002. Automated De Novo Identification of Repeat Sequence Families in Sequenced Genomes. Genome Res. 12:1269–1276. doi: 10.1101/gr.88502.

Barden LS. 1987. Invasion of *Microstegium vimineum* (Poaceae), An Exotic, Annual, Shade-Tolerant, C4 Grass, into a North Carolina Floodplain. American Midl Nat. 118:40–45. doi: 10.2307/2425626.

Baucom RS et al. 2009. Exceptional diversity, non-random distribution, and rapid evolution of retroelements in the B73 maize genome. PLoS Genet. 5:e1000732. doi: 10.1371/journal.pgen.1000732.

te Beest M et al. 2012. The more the better? The role of polyploidy in facilitating plant invasions. Ann Bot. 109:19–45. doi: 10.1093/aob/mcr277.

Bergeron M-J, Feau N, Stewart D, Tanguay P, Hamelin RC. 2019. Genome-enhanced detection and identification of fungal pathogens responsible for pine and poplar rust diseases. PLoS One. 14:e0210952. doi: 10.1371/journal.pone.0210952.

Bertolotti AC et al. 2020. The structural variation landscape in 492 Atlantic salmon genomes. Nat Commun. 11:5176. doi: 10.1038/s41467-020-18972-x.

Cantarel BL et al. 2008. MAKER: an easy-to-use annotation pipeline designed for emerging model organism genomes. Genome Res. 18:188–196. doi: 10.1101/gr.6743907.

Chan PP, Lowe TM. 2019. tRNAscan-SE: Searching for tRNA genes in genomic sequences. Methods Mol Biol. 1962:1–14. doi: 10.1007/978-1-4939-9173-0_1.

Chown SL et al. 2015. Biological invasions, climate change and genomics. Evol Appl. 8:23–46. doi: 10.1111/eva.12234.

Culpepper LZ, Wang H-H, Koralewski TE, Grant WE, Rogers WE. 2018. Understory upheaval: factors influencing Japanese stiltgrass invasion in forestlands of Tennessee, United States. Bot Stud. 59:20. doi: 10.1186/s40529-018-0236-8.

DeGiorgio M, Huber CD, Hubisz MJ, Hellmann I, Nielsen R. 2016. SweepFinder2: increased sensitivity, robustness and flexibility. Bioinformatics. 32:1895–1897. doi: 10.1093/bioinformatics/btw051.

Dobin A et al. 2013. STAR: ultrafast universal RNA-seq aligner. Bioinformatics. 29:15–21. doi: 10.1093/bioinformatics/bts635.

Emery SM, Luke Flory S, Clay K, Robb JR, Winters B. 2013. Demographic responses of the invasive annual grass *Microstegium vimineum* to prescribed fires and herbicide. For Ecol Manag. 308:207–213. doi: 10.1016/j.foreco.2013.08.002.

Flynn JM et al. 2020. RepeatModeler2 for automated genomic discovery of transposable element families. PNAS. 117:9451–9457. doi: 10.1073/pnas.1921046117.

Gibson DJ, Spyreas G, Benedict J. 2002. Life History of *Microstegium vimineum* (Poaceae), an Invasive Grass in Southern Illinois. Journal Torrey Bot Soc. 129:207–219. doi: 10.2307/3088771.

Guan D et al. 2020. Identifying and removing haplotypic duplication in primary genome assemblies. Bioinformatics. 36:2896–2898. doi: 10.1093/bioinformatics/btaa025.

Hamelin RC, Roe AD. 2020. Genomic biosurveillance of forest invasive alien enemies: A story written in code. Evol Appl. 13:95–115. doi: 10.1111/eva.12853.

van Klinken RD, Panetta FD, Coutts S, Simon BK. 2015. Learning from the past to predict the future: an historical analysis of grass invasions in northern Australia. Biol Invasions. 17:565–579 doi: 10.1007/s10530-014-0749-3.

Keriö S et al. 2020. From genomes to forest management – tackling invasive *Phytophthora* species in the era of genomics. Can J Plant Pathol. 42:1–29. doi: 10.1080/07060661.2019.1626910.

Korf I. 2004. Gene finding in novel genomes. BMC Bioinformatics. 5:59. doi: 10.1186/1471-2105-5-59.

Kreiner JM et al. 2019. Multiple modes of convergent adaptation in the spread of glyphosate-resistant *Amaranthus tuberculatus*. PNAS. 116:21076–21084. doi: 10.1073/pnas.1900870116.

Laetsch DR, Blaxter ML. 2017. BlobTools: Interrogation of genome assemblies. F1000Research. 6:1287. doi: 10.12688/f1000research.12232.1.

Larkin DJ. 2012. Lengths and correlates of lag phases in upper-Midwest plant invasions. Biol Invasions. 14:827–838. doi: 10.1007/s10530-011-0119-3.

Liu H et al. 2020. Evolution and domestication footprints uncovered from the genomes of *Coix*. Molecular plant. 13:295–308. doi: 10.1016/j.molp.2019.11.009.

Ma J, Bennetzen JL. 2004. Rapid recent growth and divergence of rice nuclear genomes. PNAS. 101:12404–12410. doi: 10.1073/pnas.0403715101.

Malinsky M, Matschiner M, Svardal H. 2021. Dsuite - Fast D-statistics and related admixture evidence from VCF files. Mol Ecol Resour. 21:584–595. doi: 10.1111/1755-0998.13265.

Manni M, Berkeley MR, Seppey M, Simão FA, Zdobnov EM. 2021. BUSCO Update: Novel and streamlined workflows along with broader and deeper phylogenetic coverage for scoring of eukaryotic, prokaryotic, and viral genomes. Mol Biol Evol. doi: 10.1093/molbev/msab199.

McCartney MA, Mallez S, Gohl DM. 2019. Genome projects in invasion biology. Conserv Genet. 20:1201–1222. doi: 10.1007/s10592-019-01224-x.

Mérel V et al. 2021. The worldwide invasion of *Drosophila suzukii* is accompanied by a large increase of transposable element load and a small number of putatively adaptive insertions. Mol Biol Evol. msab155. doi: 10.1093/molbev/msab155.

Mitros T et al. 2020. Genome biology of the paleotetraploid perennial biomass crop *Miscanthus*. Nat Commun. 11:5442. doi: 10.1038/s41467-020-18923-6.

Nees, P. 2016 *Microstegium vimineum* (Trin.) A. Camus. Bulletin OEPP/EPPO Bulletin, 46:14–19. doi: 10.1111/epp.12276.

Nielsen R et al. 2005. Genomic scans for selective sweeps using SNP data. Genome Res. 15:1566–1575. doi: 10.1101/gr.4252305.

North HL, McGaughran A, Jiggins CD. 2021. Insights into invasive species from whole-genome resequencing. Mol Ecol. doi: 10.1111/mec.15999.

Olazcuaga L et al. 2020. A whole-genome scan for association with invasion success in the fruit fly *Drosophila suzukii* using contrasts of allele frequencies corrected for population structure. Mol Biol Evol. 37:2369–2385. doi: 10.1093/molbev/msaa098.

Ou S et al. 2019. Benchmarking transposable element annotation methods for creation of a streamlined, comprehensive pipeline. Genome Biol. 20:275. doi: 10.1186/s13059-019-1905-y.

Pandit MK, Pocock MJO, Kunin WE. 2011. Ploidy influences rarity and invasiveness in plants. J Ecol. 99:1108–1115. doi: 10.1111/j.1365-2745.2011.01838.x.

Paterson AH et al. 2009. The *Sorghum bicolor* genome and the diversification of grasses. Nature. 457:551–556. doi: 10.1038/nature07723.

Paterson AH, Bowers JE, Chapman BA. 2004. Ancient polyploidization predating divergence of the cereals, and its consequences for comparative genomics. PNAS. 101:9903–9908. doi: 10.1073/pnas.0307901101.

Pejchar L, Mooney HA. 2009. Invasive species, ecosystem services and human well-being. Trends Ecol Evol. 24:497–504. doi: 10.1016/j.tree.2009.03.016.

Pimentel D, Zuniga R, Morrison D. 2005. Update on the environmental and economic costs associated with alien-invasive species in the United States. Ecol Econ. 52:273–288. doi: 10.1016/j.ecolecon.2004.10.002.

Price AL, Jones NC, Pevzner PA. 2005. De novo identification of repeat families in large genomes. Bioinformatics. 21 Suppl 1:i351–8. doi: 10.1093/bioinformatics/bti1018.

Putnam NH et al. 2016. Chromosome-scale shotgun assembly using an in vitro method for long-range linkage. Genome Res. 26:342–350. doi: 10.1101/gr.193474.115.

Qiao X et al. 2018. Different modes of gene duplication show divergent evolutionary patterns and contribute differently to the expansion of gene families involved in important fruit traits in pear (*Pyrus bretschneideri*). Front Plant Sci. 9:161. doi: 10.3389/fpls.2018.00161.

Redman DE. 1995. Distribution and habitat types for nepal *Microstegium* [*Microstegium vimineum* (trin.) Camus] in Maryland and the district of Columbia. Castanea. 60:270–275.

Schnable PS et al. 2009. The B73 maize genome: complexity, diversity, and dynamics. Science. 326:1112–1115. doi: 10.1126/science.1178534.

Schrader L et al. 2014. Transposable element islands facilitate adaptation to novel environments in an invasive species. Nat Commun. 5:5495. doi: 10.1038/ncomms6495.

Sensalari C, Maere S, Lohaus R. 2021. ksrates: positioning whole-genome duplications relative to speciation events in KS distributions. Bioinformatics. btab602. doi: 10.1093/bioinformatics/btab602.

Simão FA, Waterhouse RM, Ioannidis P, Kriventseva EV, Zdobnov EM. 2015. BUSCO: assessing genome assembly and annotation completeness with single-copy orthologs. Bioinformatics. 31:3210–3212. doi: 10.1093/bioinformatics/btv351.

Simberloff D. Invasive species: what everyone needs to know - University of Missouri-St. Louis Libraries. http://link.umsl.edu/portal/Invasive-species--what-everyone-needs-to-know/Wpr1NfXaxNo/ (Accessed August 30, 2021).

Springer NM et al. 2018. The maize W22 genome provides a foundation for functional genomics and transposon biology. Nat Genet. 50:1282–1288. doi: 10.1038/s41588-018-0158-0.

Stanke M, Diekhans M, Baertsch R, Haussler D. 2008. Using native and syntenically mapped cDNA alignments to improve de novo gene finding. Bioinformatics. 24:637–644. doi: 10.1093/bioinformatics/btn013.

Tang H et al. 2008. Unraveling ancient hexaploidy through multiply-aligned angiosperm gene maps. Genome Res. 18:1944–1954. doi: 10.1101/gr.080978.108.

The UniProt Consortium. 2021. UniProt: the universal protein knowledgebase in 2021. Nucleic Acids Res. 49:D480–D489. doi: 10.1093/nar/gkaa1100.

Tian Z et al. 2009. Do genetic recombination and gene density shape the pattern of DNA elimination in rice long terminal repeat retrotransposons? Genome Res. 19:2221–2230. doi: 10.1101/gr.083899.108.

Wang B et al. 2021. Pan-genome analysis in *Sorghum* highlights the extent of genomic variation and sugarcane aphid resistance genes. bioRxiv. 2021.01.03.424980. doi: 10.1101/2021.01.03.424980.

Wang Y et al. 2012. MCScanX: a toolkit for detection and evolutionary analysis of gene synteny and collinearity. Nucleic Acids Res. 40:e49. doi: 10.1093/nar/gkr1293.

Waterhouse RM, Zdobnov EM, Kriventseva EV. 2010. Correlating traits of gene retention, sequence divergence, duplicability and essentiality in vertebrates, arthropods, and fungi. Genome Biol Evol. 3:75–86. doi: 10.1093/gbe/evq083.

Watson L; Dallwitz MJ, 1992. The families of flowering plants: descriptions, illustrations, identification, and information retrieval. https://www.cabi.org/isc/abstract/20067201518 (Accessed September 7, 2021).

Yainna S et al. 2020. Genomic balancing selection is key to the invasive success of the fall armyworm. bioRxiv. 2020.06.17.154880. doi: 10.1101/2020.06.17.154880.

Yan Q et al. 2021. The elephant grass (*Cenchrus purpureus*) genome provides insights into anthocyanidin accumulation and fast growth. Mol Ecol Resour. 21:526–542. doi: 10.1111/1755-0998.13271.

